# An ERP index of real-time error correction within a noisy-channel framework of human communication

**DOI:** 10.1101/2020.02.08.930214

**Authors:** Rachel Ryskin, Laura Stearns, Leon Bergen, Marianna Eddy, Evelina Fedorenko, Edward Gibson

## Abstract

Recent evidence suggests that language processing is well-adapted to noise in the input (e.g., speech errors, mishearing) and readily corrects the input via rational inference over possible intended sentences and probable noise corruptions. However, it remains unclear whether this inference takes the form of an offline re-analysis or a rapid, real-time correction to the representations of the input. We hypothesize that noise inferences happen online during processing and that well-studied ERP components may serve as a useful index of this process. In particular, a reduced N400 effect and increased P600 effect appear to accompany sentences where the probability that the message was corrupted by noise exceeds the probability that it was produced intentionally and perceived accurately. Indeed, semantic violations that are attributable to noise—for example, in “The storyteller could turn any incident into an amusing <u>antidote</u>”, where the implausible word “antidote” is orthographically and phonologically close to the intended “anecdote”—elicit a reduced N400 effect and larger P600 effect. Further, the magnitude of this P600 effect is shown to relate to the probability that the comprehender will retrieve a plausible alternative. This work thus adds to the growing body of literature that suggests that many aspects of language processing are well-adapted to noise in the input and opens the door to electrophysiologic investigations of these processes

---

A recent key insight in the sentence processing literature is that the input to the comprehension system is often noisy (Levy, 2008; see also Ferreira & Patson, 2007). This noise stems from a) production errors (speech errors, typographical errors, etc.), and b) perception errors (due to sub-optimal listening/viewing conditions, noise in the environment, etc.). However, communication typically proceeds smoothly, suggesting that comprehension mechanisms are well-adapted to this noise. A rational comprehender’s guess of what was intended in a noise-corrupted linguistic exchange can be expressed as the probability of the speaker’s intended sentence, s_i_, given the perceptual input, s_p_: P(s_i_ | s_p_). By Bayes’ rule, this value is proportional to the product of the prior (what is likely to be communicated), P(s_i_), and the likelihood that a noise process would generate s_p_ from s_i_, P(s_p_ | s_i_) (e.g., Gibson et al., 2013).

Indeed, behavioral studies using offline comprehension questions suggest that readers often take the meaning of a sentence to differ from that of the literal string when that literal string has low prior probability, P(s_i_), (Ferreira, 2003), and/or the potential noise corruption that might have generated that string has high probability, P(s_p_ | s_i_) (Gibson et al., 2013). To our knowledge, only one study has explored whether this “noisy-channel” inference takes place in real time during initial sentence processing or whether it is the result of a post-interpretive process (note that these are not mutually exclusive). Levy et al. (2009) used eye-tracking to show that, when a later portion of a sentence (e.g., “The coach smiled at the player <u>tossed the ball.”</u>) renders P(s_i_) low, readers look back to previous locations in the sentence (e.g., “at”) which are probable loci of noise corruptions (e.g., because P(“at” | “as”) is high). In other words, readers maintain uncertainty about preceding input as they process a sentence and can revise their intial parse in real time.

However, the existing evidence leaves open the possibility that this uncertainty only exists with respect to linguistic material being maintained in memory (Futrell et al., 2020). For example, it is unknown whether readers additionally considered alternatives to the word that they were actively fixating (e.g., “tossed”). In the present work, we probe whether noisy-channel correction can take place in the moment of processing by leveraging the temporal resolution of the electroencephalogram (EEG) signal.

Two event-related potential (ERP) components have been consistently linked to sentence comprehension in electrophysiological investigations of language processing. The N400—a negativity peaking 400ms after word onset—is hypothesized to index the ease of semantic retrieval (e.g., after “I take my coffee with cream and…”, “dog” elicits a more negative deflection than “sugar”; Kutas & Hillyard, 1984; Kutas & Federmeier, 2011). Recent computational models construe the N400 as indexing the lexico-semantic prediction error or the update in network activation elicited by a word as it is integrated into the preceding context (e.g., Fitz & Chang, 2019; Rabovsky et al., 2018; cf. Cheyette & Plaut, 2017). Consistent with a noisy-channel view of language processing, the N400 is reduced when an incongruous completion is orthographically related to a plausible continuation, such that a plausible noise corruption might be inferred (e.g., “Before lunch he has to deposit his paycheck at the bark [vs. bank]”; Laszlo & Federmeier, 2009; Ito et al., 2016).

The P600—a positivity most pronounced 600-900ms after word onset—is less well understood. It was originally hypothesized to reflect syntactic integration difficulty (e.g., after “Every Monday he…”, “mow” elicits a larger positivity than “mows”; Osterhout & Holcomb, 1992; Friederici, 1995; Hagoort et al., 1993). However, this interpretation has faced numerous challenges. First, a number of non-syntactic manipulations elicit a P600 (e.g., spelling errors - “fone” instead of “phone”; Münte, Heinze, Matzke, Wieringa, & Johannes, 1998; van de Meerendonk, Indefrey, Chwilla, & Kolk, 2011; Vissers, Chwilla, & Kolk, 2006). Second, sentences like “The hearty meal was devouring…” elicit a P600 in spite of being syntactically well-formed (e.g., Kim & Osterhout, 2005; Kuperberg, 2007; Kuperberg et al., 2003; van Herten et al., 2005). According to traditional interpretations of these components, because these sentences are semantically anomalous, an N400 should ensue in place of these “semantic P600’s” (Brouwer et al., 2012).

Consequently, alternative accounts of the P600 have been put forward in the literature. Some appeal to parallel streams of (syntactic and semantic) processing in constructing the representation for an input string (e.g., Kim & Sikos, 2011; Kos et al., 2010; Kuperberg, 2007). Others argue that, given its scalp distribution and tight time-locking to responses, the P600 belongs to the P300 family of domain-general components (Coulson et al., 1998; Sassenhagen et al., 2014; Sassenhagen & Fiebach, 2019; for a review, see Leckey & Federmeier, 2019), which are thought to index the process of updating one’s model of the world when one encounters low-probability (“oddball”) events (Donchin, 1981; Sutton et al., 1965). Consistent with a connection to the P300, Kolk and colleagues proposed an account of the P600 as indexing our continuous monitoring of the linguistic (or other) input for possible errors (Kolk et al., 2003; Kolk & Chwilla, 2007; van de Meerendonk et al., 2011; Vissers et al., 2006). Recent computational accounts take different approaches: Brouwer et al. (2017) propose a single-stream model of N400 and P600 effects, and argue that the P600 indexes semantic integration into the unfolding utterance (conceptually similar to the N400 in Rabovsky et al.’s model). And Fitz and Chang (2019) model the P600 as the prediction error at the sequencing layer of a neural network.

Building on this error-monitoring perspective (van de Meerendonk et al., 2011), we propose that the P600, may provide a useful index of the *rational inference process* in the noisy-channel framework. When the input is anomalous but unlikely to have been an error, an N400 ensues and no P600 is typically observed. In contrast, if the input is anomalous but can be explained by a plausible noise process, readers infer that a more probable intended sentence was corrupted, and a P600 ensues while the N400 is reduced. Note that we do not aim to provide a mechanistic account of the P600 (or N400). Rather, we aim to relate well-known patterns in the EEG signal to a computational-level account of sentence comprehension (Gibson et al., 2013; Levy, 2008) in order to probe how noisy-channel inference unfolds online.

More precisely (Eq.1), given (i) a preceding sentence context and its most probable parse, *C*^1^; (ii) an expected completion word, *w*_*expected*_; (iii) the incoming (target) word: *w*_*received*_; and (iv) *s*_*expected*_ and *s*_*received*_ the sentences that correspond to connecting *w*_*expected*_ and *w*_*received*_ to *C* respectively, there is a larger P600 noise correction signal whenever P(s_i_=*s*_*received*_ | s_p_=*s*_*received*_) is lower than P(s_i_=*s*_*expected*_ | s_p_=*s*_*received*_):

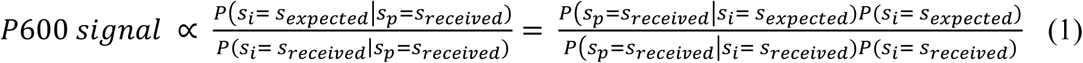

Several previously observed empirical phenomena can be reinterpreted through the lens of noisy-channel communication providing further support for the use of the P600 as an index of noisy-channel processing. First, a P600 occurs for the “traditional” syntactic violations (number/gender/case agreement errors) and for other minor deviations from the target utterance (e.g., spelling errors), because a close alternative exists in these cases, which the comprehender can correct to. For example, the probability of the meaning/structure resulting from completing “Every Monday he…” with “mow”, P(s_i_=“Every Monday he mow”), is low, while P(s_i_=“Every Monday he mows”) is relatively high. Critically, the probability of a noise process changing “mows” to “mow,” P(s_p_=“Every Monday he mow”|s_i_=“Every Monday he mows”), is relatively high; “mow*”* involves only a single character/morpheme deletion from “mows”.

Second and similarly, “semantic P600s” may manifest because a close alternative exists that the producer plausibly intended. For example, P(s_i_=“The hearty meal was devouring…”), is low, while P(s_i_= “The hearty meal was devoured…”) is relatively high, and critically P(s_p_= “The hearty meal was devouring…” | s_i_= “The hearty meal was devoured…”) is high.

Third, little to no P600 occurs for “traditional” semantic violations^2^ because the noise corruption is implausible in those cases (e.g., P(s_p_=“I take my coffee with cream and <u>dog</u>”|s_i_=“I take my coffee with cream and <u>sugar</u>”) is low).

Fourth, a much reduced or no P600 is observed for syntactic errors in “Jabberwocky” sentences, i.e., sentences that include function words/morphemes but cannot be interpreted with respect to world knowledge (Münte et al., 1997; Yamada & Neville, 2007). In such cases, it is difficult to infer plausibly intended meanings because the materials are, by design, devoid of meaning.

Finally, a P600 has been observed in studies with semantic violations in extended discourse contexts. For example, in a study by Nieuwland and Van Berkum (2005), participants read a story (e.g., about a tourist and his suitcase; both entities were mentioned several times). In critical sentences like “Next, the woman told the tourist/suitcase…”, a P600 was observed for “suitcase” (not an N400, as in a null context), plausibly because a word substitution error, when both lexical entries are highly probable in the discourse, is a probable production error. Similarly, code switches, which are probable in bilingual speech, elicit a P600 (Moreno et al., 2002).

Here, we directly evaluate whether the P600 component tracks noisy-channel inferences using an experimental design with four conditions (Table 1): (1) a control condition with no violations, (2) a condition with a canonical semantic violation, (3) a condition with a canonical syntactic violation (number agreement error), and critically, (4) a condition where the target word was semantically inappropriate but orthographically and phonologically close (e.g., in terms of Levenshtein distance) to a semantically plausible neighbor. Behavioral norming indicates that the proximity of such a neighbor makes the plausibly intended word recoverable. As a result, the critical condition is expected to elicit a noisy-channel inference and, by hypothesis, a P600 (similar to the syntactic condition and in contrast to the semantic condition).

**Table 1.** Example materials and predictions.

| **The storyteller could turn any incident into an amusing ...** |  |  |  |
| --- | --- | --- | --- |
| **Completion** | **Condition** | **N400 prediction** | **P600 prediction** |
| anecdote | Control | - | - |
| hearse | Semantic violation | + | - |
| anecdotes | Syntactic violation | - | + |
| antidote | Critical | +/- | + |

## Methods

### Participants

Twenty-nine right-handed native English speakers participated in this study, 24 of whom were included in the final analysis (10 males; age 18-40 years). Participants were recruited from the MIT Brain and Cognitive Sciences subject pool and the Wellesley College student community. Informed consent was obtained in accordance with the MIT Committee on the Use of Humans as Experimental Subjects. Participants were compensated with cash for their participation. Five subjects were excluded from final analysis due to an excessive number of artifacts in the EEG signal.

### Materials

One hundred sixty ten-word-long sentences were constructed (with 4 conditions each, as described above) and distributed across four presentation lists following a Latin Square design, so that each list contained only one version of an item (and 40 trials per condition). The target word (always a noun) was the last word in the sentence. The target words in the semantic violation and critical conditions were target words in the control condition for other items (e.g., “hearse” in the example above was the target word in the control condition of another item); the target words were thus identical across these conditions (and were only different in the number feature between these conditions and the syntactic violation condition). Materials were normed on an independent set of participants to ensure that a) the target words were judged less likely to be errors in the semantic violation condition than in the critical and syntactic violation conditions, and b) the intended words were more recoverable in the critical and syntactic violation conditions than in the semantic violation condition (see OSF repository for details: https://osf.io/vcsfb/?view_only=ba0079719cfa4118be5cc99714135acf). In addition, 320 10-word-long filler items were constructed. These contained no semantic or syntactic violations.

### EEG recording

EEG was recorded from 32 scalp sites (10-20 system positioning), a vertical eye channel for detecting blinks, a horizontal eye channel to monitor for saccades, and two additional electrodes affixed to the skin above the mastoid bone. EEG was acquired with the Active Two Biosemi system using active Ag-AgCl electrodes mounted on an elastic cap (Electro-Cap Inc.). All channels were referenced offline to an average of the mastoids. The EEG was recorded at 512 Hz sampling rate and filtered offline (bandpass 0.1-40 Hz). Trials with blinks, eye movements, muscle artifact, and skin potentials were rejected prior to averaging and analysis. An average of 15.6% of trials were rejected per participant (range: min = 0.6%, max = 26.3%).

### Testing procedure

Participants were tested individually in a sound-attenuated booth where stimuli were presented on a computer monitor. Stimuli appeared in the center of the screen word-by-word, time-locked to the vertical refresh rate of the monitor (75 Hz). The sentences were displayed word-by-word in white on a black background. Each trial began with a pre-trial fixation (1,000 ms), followed by 500 ms of a blank screen. Then, the sentence was presented for 5,800 ms (400 ms per word and 100 ms ISI, with an ISI of 900 ms after the last word). The order of trials was randomized separately for each participant. Each list was pseudo-randomly divided into ten “runs”, in order to give participants breaks as needed. Each run contained 4 trials of each condition and 32 fillers.

To ensure that participants read the sentences for meaning, yes/no comprehension questions appeared after 60 of the 480 trials (experimental and filler), constrained such that there were no more than three consecutive trials with a question, and no more than 20 consecutive trials without a question. The correct answer was “yes” half of the time. Comprehension questions were displayed all at once (for 3,500 ms + 100 ISI) in aqua on a black background, and participants responded “yes” or “no” by pressing buttons on a gamepad. At the beginning of the experiment, participants were shown a set of 12 practice items to familiarize them with the procedure. The experiment took ∼1 hour.

### Analysis

Eight centro-parietal electrode sites (C3, Cz, C4, CP1, CP2, P3, Pz, and P4) were included in the analysis. These sites reflect the typical distribution of N400 and P600 effects reported in the literature (Kutas & Federmeier, 2011; Tanner, 2019). ERP signals were time-locked to the onset of the sentence-final (target) word and individual trial epochs from 100 ms prior to the onset of this stimulus until 1,000 ms after onset were extracted. The time window from -100 ms to word onset was used as the pre-stimulus baseline. Mean amplitude measurements were computed in two time windows – 300-500 ms and 600-800 ms – to quantify the N400 and P600 components, respectively. Time windows were chosen to match standard time windows used in the literature (Gouvea et al., 2010; Kutas & Federmeier, 2011) and to be equal in duration with a 100 ms gap in between to reduce dependence between the windows.

For each of the two time windows of interest (300-500 ms and 600-800 ms), the mean amplitude was entered as the dependent variable in a linear mixed-effects regression model, with condition (control, semantic violation, syntactic violation, critical) as a dummy-coded fixed effect (with control as the reference level). The models included random intercepts for participants, items, and electrodes, and random condition slopes for each grouping variable. Analyses were performed using the “brms” package for Bayesian regression modeling in R (Bürkner, 2017), which interfaces with the Stan probabilistic programming language (Carpenter et al., 2017). Moderately regularizing priors were chosen based on prior literature. In particular, a normal distribution with mean 0 and standard deviation 2.5 was chosen for the beta coefficients based on the reasoning that an ERP effect of +/- 5μV is fairly common. Data and analysis code are available at https://osf.io/vcsfb/?view_only=ba0079719cfa4118be5cc99714135acf.

## Results

Participants mostly answered the comprehension questions accurately (mean = 0.88, bootstrapped 95% confidence interval = [0.85, 0.91), which suggests that they were engaged in the task.

### N400 and P600 components

As expected, and replicating many previous studies, in the N400 window, the ERP amplitude decreased by -4.09 μV (95% Credible Interval (CI) = [-5.06, -3.02]) in the semantic condition relative to the control condition. The amplitude was also somewhat more negative (Estimate = -1.37, 95% CI = [-2.51, -0.17]) in the critical condition relative to the control condition but not in the syntactic condition (Estimate = -0.48, 95% CI = [-1.66, 0.72]). (An N400 effect is expected for the critical condition target word because it is not strongly facilitated by the semantic context, unlike the control condition target word.) In the P600 window, the ERP amplitude did not differ between the control condition and the semantic condition (Estimate = -0.85, 95% CI = [-2.08, 0.35]).

However, P600 amplitude was more positive both in the syntactic (Estimate = 2.10, 95% CI = [0.91, 3.22]) and in the critical condition (Estimate = 1.34, 95% CI = [0.11, 2.52]). In other words, as predicted by the noisy-channel inference account, the critical condition, where the target word was semantically inappropriate but phonologically and orthographically close to a plausible neighbor, elicited a P600 effect, similar to the syntactic condition. See Figures 1 and 2 for summaries and https://osf.io/vcsfb/?view_only=ba0079719cfa4118be5cc99714135acf for full model estimates.

**Figure 1.**
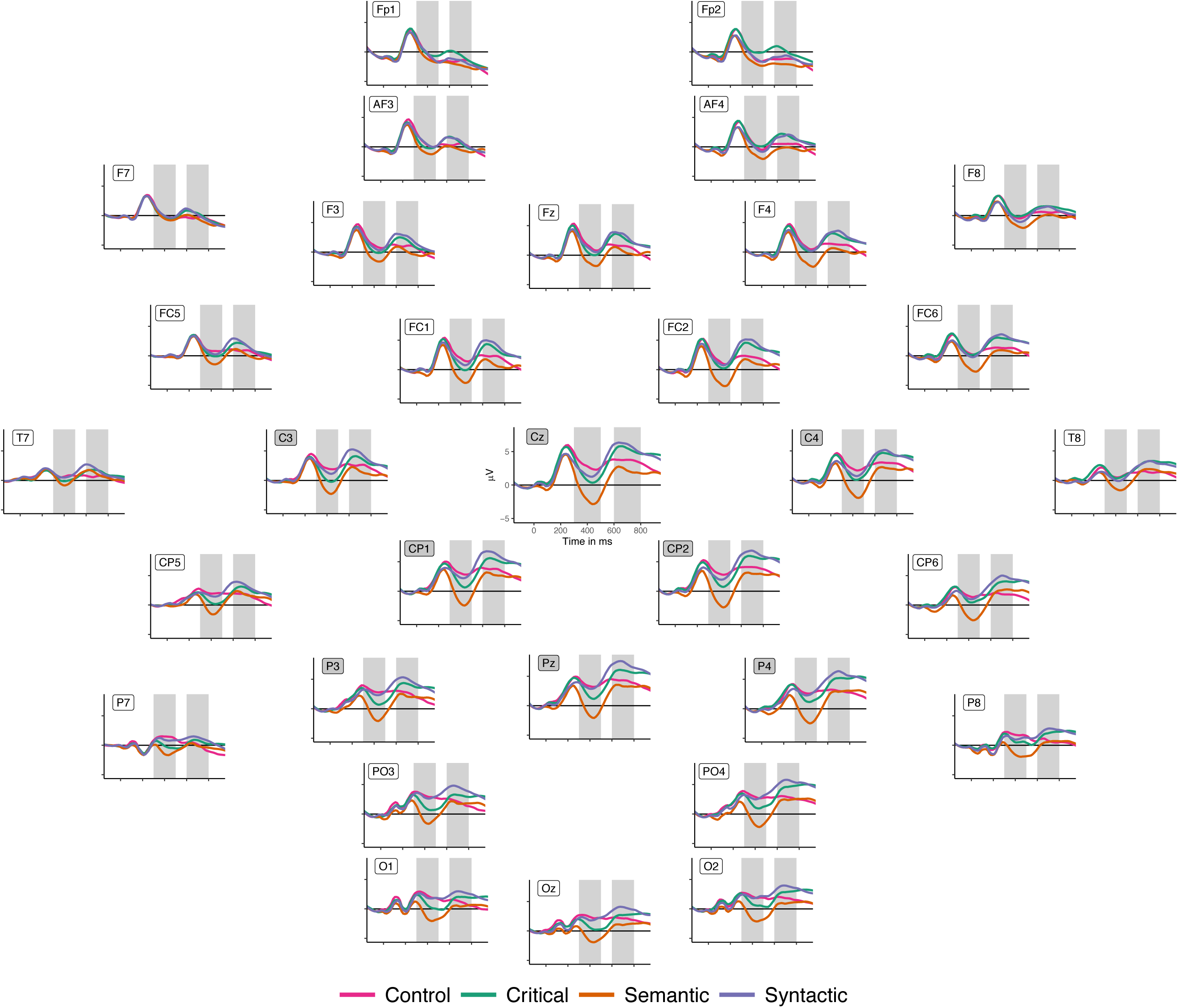
Grand average ERPs for each condition at every recorded electrode. The x-axis shows time from the onset of the presentation of the final word, and the y-axis shows voltage (negative plotted down), as compared to the mean voltage of the baseline 100 ms pre-stimulus interval. (The subset of channels used in the statistical analyses is indicated by the gray labels and the two gray rectangles in each plot indicate the time windows of interest: 300-500ms and 600-800ms.)

**Figure 2.**
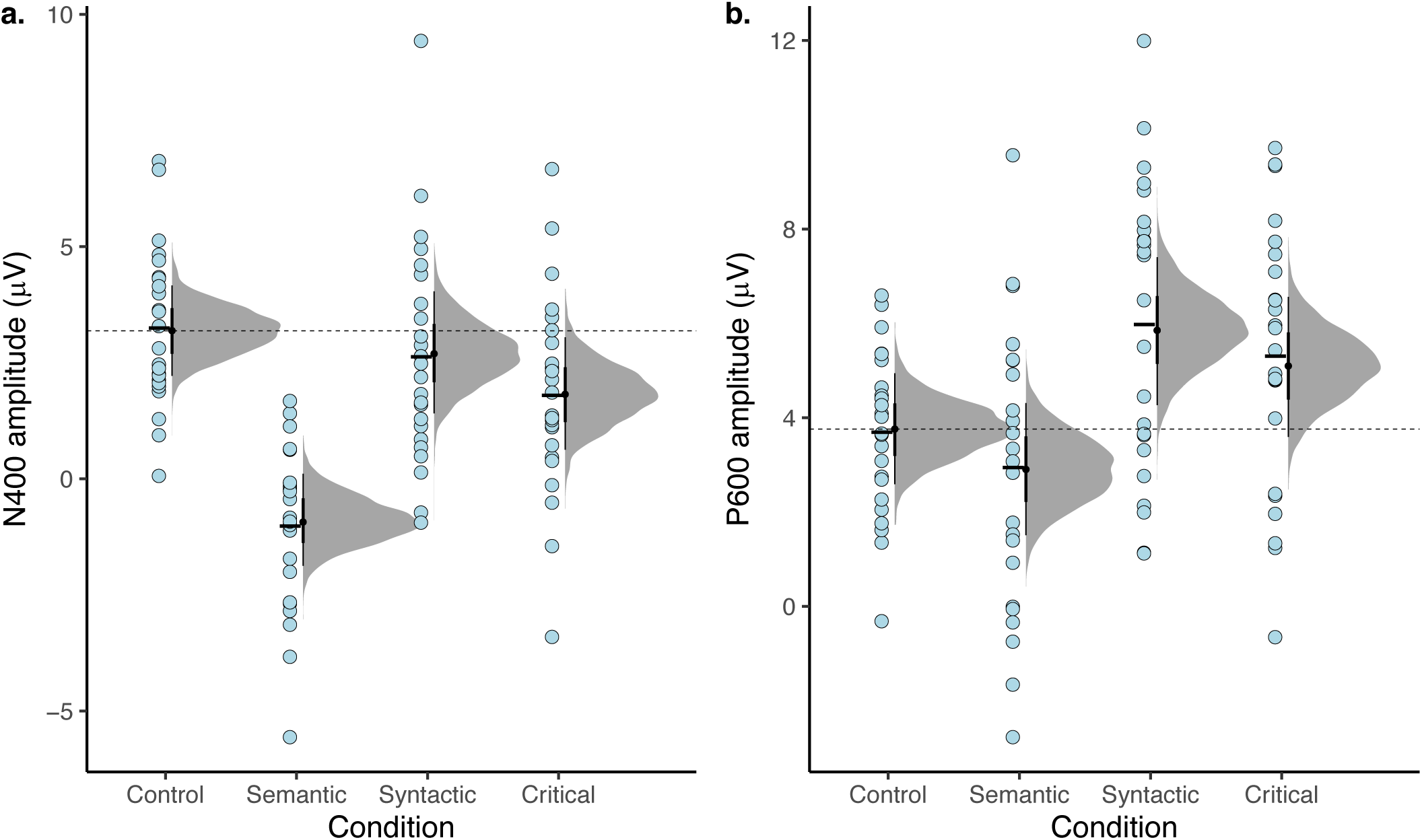
Mean amplitudes of (a) the N400 and (b) P600 components. Light blue points represent individual participant means and the black horizontal bar represents the overall mean for each condition. Densities and point intervals represent the distribution of fitted conditional means from Bayesian linear mixed-effects model posteriors. Dashed line indicates the mean amplitude in the control condition.

### Magnitude of P600 and recoverability of the plausible alternative

To further explore these effects, we assessed whether the magnitude of the P600 is linearly related to the recoverability of the word. We computed two measures of recoverability. The first is the Levenshtein distance between each target word (e.g., antidote) and its control condition counterpart (e.g., anecdote). Levenshtein distance was computed using the adist() function in R. The second measure was taken from the norming data (summary available at https://osf.io/vcsfb/?view_only=ba0079719cfa4118be5cc99714135acf): the percentage of correct guesses about which word was intended. The relationships between the magnitude of the P600 effect for an item (averaging over participants and electrodes and subtracting the P600 amplitude for the control condition from the amplitudes in the other three conditions) and the two measures of recoverability are shown in Figure 3. Three simple linear regression models were fitted using brms, with the same priors as in the above models where applicable (see further analysis details at https://osf.io/vcsfb/?view_only=ba0079719cfa4118be5cc99714135acf). Items with a larger Levenshtein distance from their control version were less likely to elicit successful recovery of the control version (Estimate = -6.93, 95% CI = [-7.54, -6.34]), confirming the validity of operationalizing recoverability as Levenshtein distance from the nearest neighbor. Items with a larger Levenshtein distance from their control also elicited smaller P600 effects (Estimate = -0.42, 95% CI = [-0.58,-0.26]). Similarly, items for which participants were more likely to recover the control elicited larger P600 effects (Estimate = 3.29, 95% CI = [1.76, 4.85]). Note that these bivariate relationships are somewhat expected given that the 3 conditions were designed to be differentially recoverable. Models which include condition as an additional covariate indicate that these two predictors (condition and Levenshtein distance or Percent recovered) explain largely redundant variance (i.e., neither predictor is estimated to have a non-zero independent contribution).

**Figure 3.**
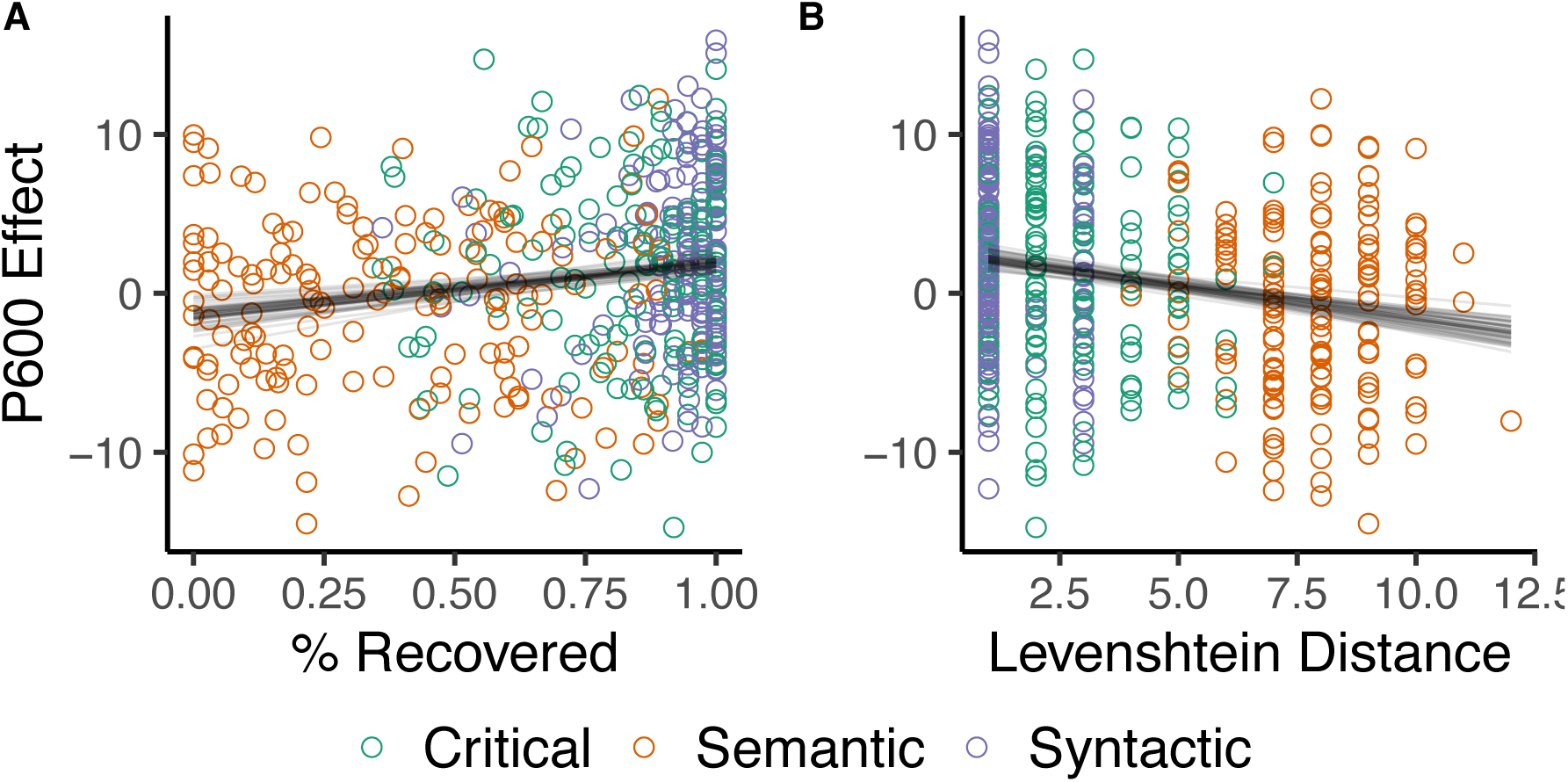
Relationships between the average P600 effect for each item in each experimental condition (after subtraction of P600 amplitude in the Control condition) and two measures of recoverability: Percent of correctly recovered completions (A) and Levenshtein distance (B). Gray lines represent 50 fitted regression lines (randomly sampled from model posteriors).

## Discussion

We observed a P600 when participants read sentences where the target word was semantically inappropriate but had an appropriate orthographic and phonological neighbor, allowing for the possibility that the received message was corrupted by noise. The intended (plausible) word was thus recoverable, and comprehenders could correct the signal. This effect was similar to that observed for the canonical syntactic violation condition. No P600 was observed for the canonical semantic violation, where the intended meaning could not be recovered. Thus, in addition to considering that memory for earlier parts of a sentence might be noisy (Futrell et al., 2020; Levy et al., 2009), readers can also correct a word to a more plausible alternative, in the moment of processing. Further, the size of the P600 was linearly related to the likelihood of recovering the plausible alternative, suggesting that this component could be leveraged to probe the reader’s implicit inferences about the noise process.

A key prediction of this proposal is that the P600 should be modulated by the distribution of errors in the input because a rational comprehender will tune their noise model to the observed distribution of errors in the environment (Gibson et al., 2013; Ryskin et al., 2018). Indeed, increasing the number of sentences that contain syntactic violations leads to a reduction of the P600 magnitude (Coulson et al., 1998; Hahne & Friederici, 1999). Similarly, Hanulíková et al. (2012) showed reduced P600s to syntactic errors in foreign-accented speech, where an agreement error is more expected (compared to native-sounding speech), suggesting that listeners take speaker-specific information into account, in addition to the overall proportion of errors in the input. Future work is needed to provide a systematic test of the quantity and nature of input that will shift the noise model and, consequently, the P600.

The current proposal does not aim to provide a mechanistic model of how the P600 is generated but rather build a bridge between the growing noisy-channel literature and the wealth of psycholinguistic studies using ERPs. Recent models of the P600 (e.g., Brouwer et al., 2017; Fitz & Chang, 2019) have focused on addressing how the comprehension mechanisms compute sentence meanings or learn their relative probabilities. The noisy-channel framework concerns the *probability* of an intended sentence given the perceived form, no matter how the most likely representations for an input string might have been computed. Thus, to the extent that they generate accurate predictions regarding the probability of perceived sentences and their alternatives, these models are compatible with the noisy-channel framework. One consideration is that these models are typically trained on clean, error-free data. Given the evidence of robustness of the human comprehension system to noise, it would be interesting to explore whether models may improve their fit to human data by training on data with (plausible) noise.

A potential limitation of this proposal is that there exist examples of P600 effects which cannot, at first glance, be readily tied to a noisy-channel correction process. Though it is beyond the scope of this paper to review the entirety of the P600 literature, we discuss several of these examples here in hopes that this will spur future investigation. As mentioned above, canonical N400s effect are sometimes followed by P600s. The addition of an explicit task (e.g., grammaticality judgment) may provide a partial explanation (but see Van Petten & Luka, 2012) – the task may increase the likelihood of noise across the board (Gibson et al., 2013). More importantly, for words that are low probability in context it will often be possible to treat it as either a faithfully represented word that is unexpected, leading to an N400, or a corrupted version of an expected word, leading to a P600-like response. Different participants may have different relative weightings of the two options given their own idiosyncratic language experience, so a blended response could reflect averaging across participants. More intriguingly, individual participants may have high uncertainty about these items and this could be reflected in diminished N400 and P600 magnitudes within the same individual. Finally, the majority of semantic violation tasks were not designed with noise-correction questions in mind and thus are unlikely to have thoroughly controlled how likely the materials are to have been corrected to an alternative, as we have done here. A re-analysis of existing datasets showing biphasic N400/P600 responses with this variable in mind may be a fruitful avenue for future progress on this topic.

In addition, pragmatic processing has been linked to P600 effects (see Hoeks & Brouwer, 2014 for a review), for example in experiments looking at comprehension of figurative language (Regel et al., 2011) and jokes (Du et al., 2013). It is possible that jokes, for instance, violate the reader’s expectation (i.e., their literal meaning has lower prior probability) and lead them to consider the alternative that would have been said, if the sentence were meant to be serious. We leave it to future work to investigate this intriguing speculation. Critically, this puzzle doesn’t undermine the utility of the P600 as an index of noisy-channel correction in future experiments, provided experimenters are careful to control pragmatics in their materials.

## Conclusion

To conclude, the P600 ERP component is promising as a signal of noisy-channel error correction taking place in real-time. Though future studies are needed to generalize this finding to a wider set of scenarios, a P600 is predicted whenever the received input can be explained as a perceptual or production error. This work contributes to a growing literature suggesting that the human language system is well-adapted to potential corruption of the linguistic signal and opens the door to investigation of the comprehender’s implicit noise model.

## Acknowledgments

We thank Roger Levy, Peter Hagoort, Gina Kuperberg, and the audiences at the Neurobiology of Language 2012 conference, and the CUNY 2013 Sentence Processing conference for feedback on this work. We are also grateful to Steve Piantadosi for comments on a draft of the manuscript. This work was supported by National Science Foundation Grant 0844472 from the Linguistics Program to EG, by the K99/R00 grant 057522 from NICHD to EF, and F32 015163 from NIDCD to RR.

## Footnotes

1 For the current purposes, we set aside the possibility of multiple parallel parses of the preceding context, *C*, and how their relative probabilities can be re-weighted given new input but see Levy et al. (2009) for discussion.

2 In some studies, a P600 is reported after an N400 for canonical semantic violations (see Brouwer, Fitz, & Hoeks, 2012; Van Petten & Luka, 2012). It is noteworthy that this is more likely to occur when an unnatural secondary task (e.g., acceptability judgments) is included. Kolk et al. (2003) directly compared judgment and comprehension tasks and found a P600 in the semantic violation cases only for the former. When the task is to find errors, participants plausibly increase the likelihood of errors across the board. Other aspects of the task, e.g., the proportion of errors in the fillers or the proportion of incongruous sentences in the environment, also affect the prior and likelihood and, therefore, the probability of the P600 on the current account (see also Delaney-Busch et al., 2019). We return to this issue in the discussion.

